# RLBWT-Based LCP Computation in Compressed Space for Terabase-Scale Pangenome Analysis

**DOI:** 10.64898/2026.01.23.701410

**Authors:** Ahsan Sanaullah, Nathaniel K. Brown, Pramesh Shakya, Arun Deegutla, Ardalan Naseri, Ben Langmead, Degui Zhi, Shaojie Zhang

## Abstract

Lossless full text indexes are utilized in a myriad of applications in bioinformatics. The continuously decreasing cost of generating biological data has resulted in the need to build full text indexes on biological datasets of increasing size. Many compressed full text indexes have been developed to address this problem. In particular, run-length Burrows-Wheeler transform (RLBWT) based compressed full text indexes have seen wide development and adoption. However, the construction of these RLBWT-based compressed full text indexes is still computationally expensive, sometimes prohibitively so, even for current dataset sizes. Therefore, we present algorithms for the construction of RLBWT-based compressed full text indexes and their supporting data structures in compressed space. The algorithms have a space complexity of *O*(*r*) words and run in *O*(*n*) time for repetitive datasets, where *r* is the number of runs in the BWT, *n* is the length of the text, and repetitive datasets implies the average run length is at least log *n*. We provide the first algorithm to compute LCP-related information for repetitive datasets in optimal time and *O*(*r*) space, greatly reducing memory requirements. The key idea behind this algorithm is the utilization of *r* samples of the inverse suffix array at regular intervals. For example, on the Human Pangenome Reference Consortium Release 2 dataset, this reduces peak memory from 2,135 GiB to 170 GiB (12.6x reduction) compared to the previous best method (pfp-thresholds).

**Availability:** The implementation is available at https://github.com/ucfcbb/TeraTools.

**Supplementary Information:** Supplementary Material is available online at bioRxiv.

## 1 Introduction

Full text indexes have been used to efficiently solve many bioinformatics problems including whole genome alignment [Delcher et al., 1999, Lin and Hsu, 2020], read mapping [Langmead et al., 2009, Li and Durbin, 2009], read alignment [Li and Durbin, 2010], identical by descent (IBD) segment detection [Freyman et al., 2020, Naseri et al., 2019], metagenomic or taxonomic classification of sequences [Kim et al., 2016], haplotype imputation and phasing [Delaneau et al., 2019, Rubinacci et al., 2020, Sanaullah et al., 2023], and variant calling [Prezza et al., 2020]. Indexes that have been used and/or developed to solve these problems include suffix trees, suffix arrays, and Burrows–Wheeler transform based indexes. The utility of solving these bioinformatics problems incentivized the generation of more biological data and the development of better, cheaper, faster biological data production methods.

As DNA sequencing data becomes cheaper and easier to generate, genomic datasets and problems are getting exponentially larger. Fortunately, the datasets also typically get more repetitive. Large-scale sequencing initiatives such as the Human Pangenome Reference Consortium [Liao et al., 2023], the All of Us Research Program [The All of Us Research Program Genomics Investigators, 2024], the UK Biobank [The UK Biobank Whole-Genome Sequencing Consortium, 2025], the Darwin Tree of Life Project [The Darwin Tree of Life Project Consortium, 2022], and the Earth BioGenome Project [Lewin et al., 2022] are currently generating petabytes of data. At this scale, traditional full text indexes that use memory proportional to the size of the dataset become too hard to build and use.

Compressed full text indexes have emerged as the key to enabling fast substring queries over large genomic collections such as pangenomes. Remarkably, these indexes support query times comparable to those of uncompressed full-text indexes while only requiring space proportional to the non-redundant content of the data. As a result, when additional (and often similar) biological sequences are incorporated, the index size grows sublinearly while still supporting fast query times. Among these indexes that achieve space usage proportional to the non-redundant content of the data while supporting exact pattern matching in time nearly linear in the query length, compressed full-text indexes based on the run-length-compressed Burrows–Wheeler Transform (RLBWT) and its variants have proven particularly effective [Bertram et al., 2024, Nishimoto and Tabei, 2021, Rossi et al., 2022, Shakya et al., 2025, Zakeri et al., 2024].

Despite these recent advances in compressed full-text indexing, constructing these indexes efficiently remains challenging. Besides the RLBWT, many practical indexes rely on related structures such as the suffix array (SA) or longest common prefix (LCP) array, which are still difficult to compute over large genomic datasets. The primary bottleneck is peak memory consumption. LCP-related structures are particularly memory intensive to compute. A recent survey [Fischer and Ohlebusch, 2025] on LCP construction states that the current best practical algorithm is a prefix-free-parsing (PFP) [Boucher et al., 2019] based method, pfp-thresholds [Rossi et al., 2022]. However, in practice, PFP based methods do not scale well to very large datasets. For example, pfp-thresholds requires about 2 days and 2.1 terabytes of RAM to compute thresholds (an LCP-related structure) for the Human Pangenome Reference Consortium (HPRC) version 2 release comprising 466 human haplotypes (run on a x2iedn.24xlarge compute instance from Amazon Web Services with 96 3.5 Ghz cores and 3 terabytes of RAM). These resource requirements are significant roadblocks for constructing compressed full-text indexes of large datasets.

Therefore, there is a strong need for algorithms that run efficiently (ideally achieving the theoretically optimal *O*(*n*)-time and *O*(*r*)-space) when building the full suite of components making up RLBWT-based compressed full text indexes, such as move structures [Nishimoto and Tabei, 2021] and *O*(*r*) samples of the SA or LCP array needed for *r*-index construction [Gagie et al., 2020]. In particular, we lack efficient methods for constructing “text-space” mappings like *ϕ* and *ϕ*^−1^, or for computing LCP summaries directly from run-length compressed inputs.

All current methods for the construction of supporting structures (i.e., SA and LCP related) of RLBWT-based compressed full text indexes either use *ω*(*r*) memory or don’t compute all supporting structures. The only tools known to the authors that compute LCP-related information in compressed space are rlbwt2lcp [Prezza and Rosone, 2021] and pfp-thresholds [Rossi et al., 2022]. rlbwt2lcp uses *O*(*n* log *σ*) bits 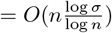 words of memory, while pfp-thresholds uses memory proportional to |*PFP* |, the sum of the sizes of the dictionary and parse of a prefix-free-parse of the text. It has been shown that |*PFP* | is asymptotically larger than 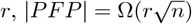), and that |*PFP*| *> r* empirically for many repetitive texts [Nishimoto et al., 2022]. Memory-efficient construction methods for RLBWT support structures currently exist neither in theory (in *O*(*r*) space) nor in practice (feasible for the largest current datasets). However, memory efficient construction methods for the RLBWT itself do exist, both in theory (*O*(*r*) words [Li, 2024, Nishimoto et al., 2022]) and in practice [Díaz-Domínguez and Navarro, 2023, Li, 2024, Olbrich, 2025]. See Supplementary Table S1 for a comparison of state of the art compressed space RLBWT construction algorithms. Also see Table 1 for a comparison of compressed space RLBWT support structure construction algorithms. Note that the exact problem we solve in this paper, time and memory efficient construction of RLBWT support structures, was recently proposed as an important problem for future work in a survey on computing very large BWTs by Díaz-Domínguez et al. [2025]. Besides ropebwt3 [Li, 2024], which doesn’t compute any LCP information and has worse time complexities than our algorithms (*O*(*n* log *r*) vs. *O*(*n* + *r* log *r*)), the algorithms presented in this paper are the only RLBWT support structure construction algorithms to achieve *O*(*r*) words of space.

**Table 1:**
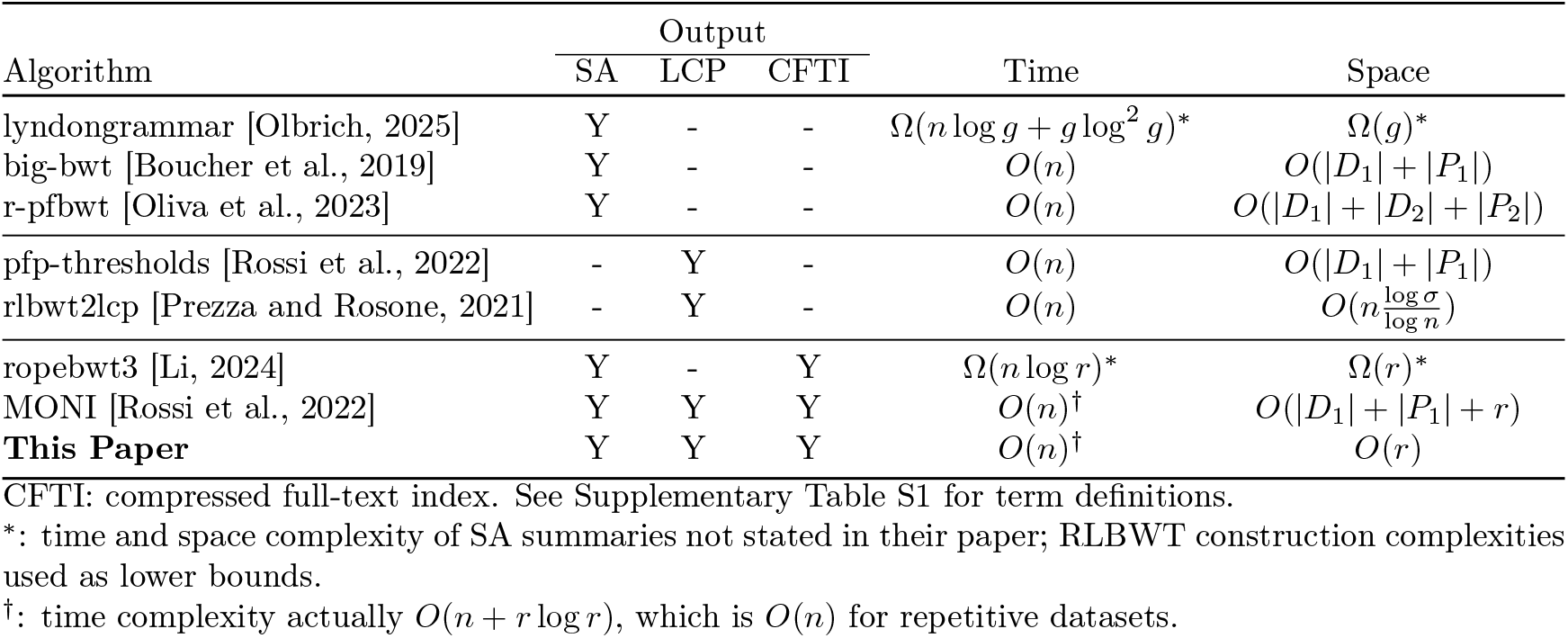
RLBWT Support Structure Construction Algorithms.

Since memory efficient RLBWT construction algorithms already exist, in this paper we present efficient compressed full-text index construction algorithms that take the RLBWT as input. Given the RLBWT of the text, our methods compute all other components of the index—including the irreducible LCP values and move structures for *ϕ, ϕ*^−1^, *LF*, and *ψ*—in overall *O*(*n*) time and *O*(*r*) space for repetitive texts (texts where 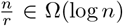). As a direct result, we obtain *O*(*n*) time and *O*(*r*) words of space algorithms for outputting all supporting structures produced by other compressed memory algorithms, including various suffix array and LCP array sampling schemes (e.g., at the beginning and ends of runs, evenly spaced samples, minimum LCP per run, and thresholds: position of minimum LCP between runs of equal characters). At the core of our approach is a novel *O*(*n*)-time, *O*(*r*)-space algorithm for computing the irreducible PLCP values from a sample of the inverse suffix array, which in turn enables the computation of LCP thresholds and similar summaries needed for advanced matching-statistics and maximal exact match queries. The algorithms presented here enable the computation of LCP summaries on large datasets such as human472 (a superset of HPRC release 2 [Liao et al., 2023]) and CommonBacteria (almost all of AllTheBacteria [Hunt et al., 2025]). To our knowledge, no other tool has generated LCP summaries of CommonBacteria, and the only tool that has computed a compressed full-text index of it is ropebwt3 [Li, 2024]. Note that preempting our algorithms with r-comp [Nishimoto et al., 2022] results in an *O*(*r*) words of space *O*(*n*) time algorithm for constructing the RLBWT, RLBWT-based compressed full-text indexes, and all RLBWT support structures of repetitive texts (texts where 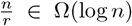). These results advance the construction and usability of run-length-compressed full-text indexes, enabling scalable, high-performance querying across modern pangenome-scale datasets.

## 2 Background

A string *S* is a sequence of characters *S*[1]*S*[2]*S*[3] … *S*[*m*] where *S*[*i*] ∈ Σ for all *i* and Σ = {1, 2, 3, …, *σ}*. The alphabet of *S* (Σ) is ordered and is said to have size *σ*. The length of *S*, is represented by |*S*| and is the number of characters it contains, |*S*| = *m*. The *i*-th character of *S* is *S*[*i*]. Suffix *i* of *S* is the substring that starts at character *i* and ends at the end of the string: *S*[*i* … *m*]. The empty string, *ε*, is the string of length 0. Suffix *m* + 1 of *S* is *ε*. A string *S* is lexicographically smaller than a string *T* (*S* ≺ *T*) if *S* = *ε* ≠ *T* or *S*[1] *< T* [1] or *S*[1] = *T* [1] and *S*[2 … *m*] ≺ *T*[2 … |*T*|]. Prefix *i* of *S* is the substring starting at the beginning of *S* and ending at *i*: *S*[1 … *i*]. The longest common prefix of strings *S* and *T, lcp*(*S, T*), is the largest string that is a prefix of both *S* and *T*.

### 2.1 Interval Sequences and Move Data Structures

A permutation *f* : [1, *n*] → [1, *n*] is composed of *r* “input intervals” [*i, j*] s.t. *f* (*i*^′^) − *i*^′^ = *f* (*i*) − *i* for all *i*^′^ ∈ [*i, j*]. A representation of such a permutation in *O*(*r*) space is a disjoint interval sequence, *I*_*f*_. *I*_*f*_ stores *r* pairs of integers (*p*_*i*_, *f* (*p*_*i*_)) where *p*_*i*_ is *i*-th smallest starting position of an input interval. Then given *j* and *x* (*j* an index in [1, *n*] and *x* ∈ [1, *r*] the index of the input interval containing *j*), where *x* = max {*k*|*p*_*k*_ ≤ *j}, f* (*j*) can be computed in constant time by *f* (*j*) = *f* (*p*_*x*_) + *j* − *p*_*x*_.

However, in order to compose *f*, i.e. to compute *f* (*f* (*j*)), one needs to compute *x*^′^ as well, where *x*^′^ is the index of the input interval containing *f* (*j*): *x*^′^ = max {*k*|*p*_*k*_ ≤ *f* (*j*)}. Nishimoto and Tabei [2021] show that the *r* intervals of *f* can be split into at most 2*r* intervals s.t. *x*^′^ can be computed in constant time and *O*(*r*) space. The result is called the balanced interval sequence, for which Bertram et al. [2024] describe an *O*(*r* log *r*) time and *O*(*r*) space construction algorithm. The data structure that computes *x*^′^ and *f* (*j*) in *O*(*r*) space and constant time given *j* and *x* is called the move data structure (call it ℳ_*f*_). The query that computes *x*^′^ and *f* (*j*) given *j* and *x* is called the move query. ℳ_*f*_ (*j, x*) = *Move*(*j, x*) = (*f* (*j*), *x*^′^).

### 2.2 Run-Length Compressed Burrows-Wheeler Transform

In the context of the Burrows-Wheeler Transform (BWT, also sometimes referred to as *L*, for “last” column of the Burrows-Wheeler matrix), strings end in a terminator character (denoted $) that doesn’t appear anywhere else in the string and is lexicographically smaller than all other characters. Here we consider the Burrows-Wheeler Transform of an ordered set of strings {*S*_1_, *S*_2_, …, *S*_*N*_*}* to be the BWT of their concatenation separated by unique termination characters (*S*_1_$_1_*S*_2_$_2_ … *S*_*N*_ $_*N*_). We call the concatenated string the “text,” *T*. It has length |*T*| = *n*, and alphabet size *σ*. $_*i*_ *<* $_*j*_ if *i < j* and $_*i*_ *<* $ and $_*i*_ occurs exactly once for all *i*. Therefore the first *N* characters of the alphabet are the sentinel characters, character *i* = $_*i*_. The suffix array, *SA*, of a text *T* is the ordering of the suffixes of the string from lexicographically smallest to largest. Suffix *SA*[*i*] of *T* (i.e. *T* [*SA*[*i*] … *n*]) is its *i*-th lexicographically smallest suffix. Since $_*i*_ has exactly one occurrence in the text and is the *i*-th lexicographically smallest character in *T, T* [*SA*[*i*]] = $_*i*_ for *i* ∈ [1, *N*].

The *i*-th character of the BWT of a text *T* is the character that precedes its *i*-th lexicographically smallest suffix. Therefore, *BWT* [*i*] = *T* [*SA*[*i*] − 1]. There are a few data structures related to the BWT and SA: LF, *ϕ*, LCP, and PLCP. LF maps from a location in SA to the location of the preceding suffix of *T* in the *SA*: *SA*[*LF* [*i*]] = *SA*[*i*] − 1. *ϕ* stores the suffix that precedes a suffix in the suffix array: *ϕ*[*SA*[*i*]] = *SA*[*i* − 1]. LCP stores the length of the longest common prefix between the *i*-th lexicographically smallest suffix and the *i* − 1-th lexicographically smallest suffix: *LCP* [*i*] = |*lcp*(*T* [*SA*[*i*] … *n*], *T* [*SA*[*i* − 1] … *n*])|. PLCP stores the length of the longest common prefix between suffix *i* and the lexicographically biggest suffix lexicographically smaller than suffix *i*: *PLCP* [*SA*[*i*]] = *LCP* [*i*]. *ISA* is the inverse of *SA, ψ* is the inverse of *LF*, and *ϕ*^−1^ is the inverse of *ϕ*. These permutations are depicted for *T* = *aacaaatttaaagagattt*$ in Figure 1.

**Figure 1.**
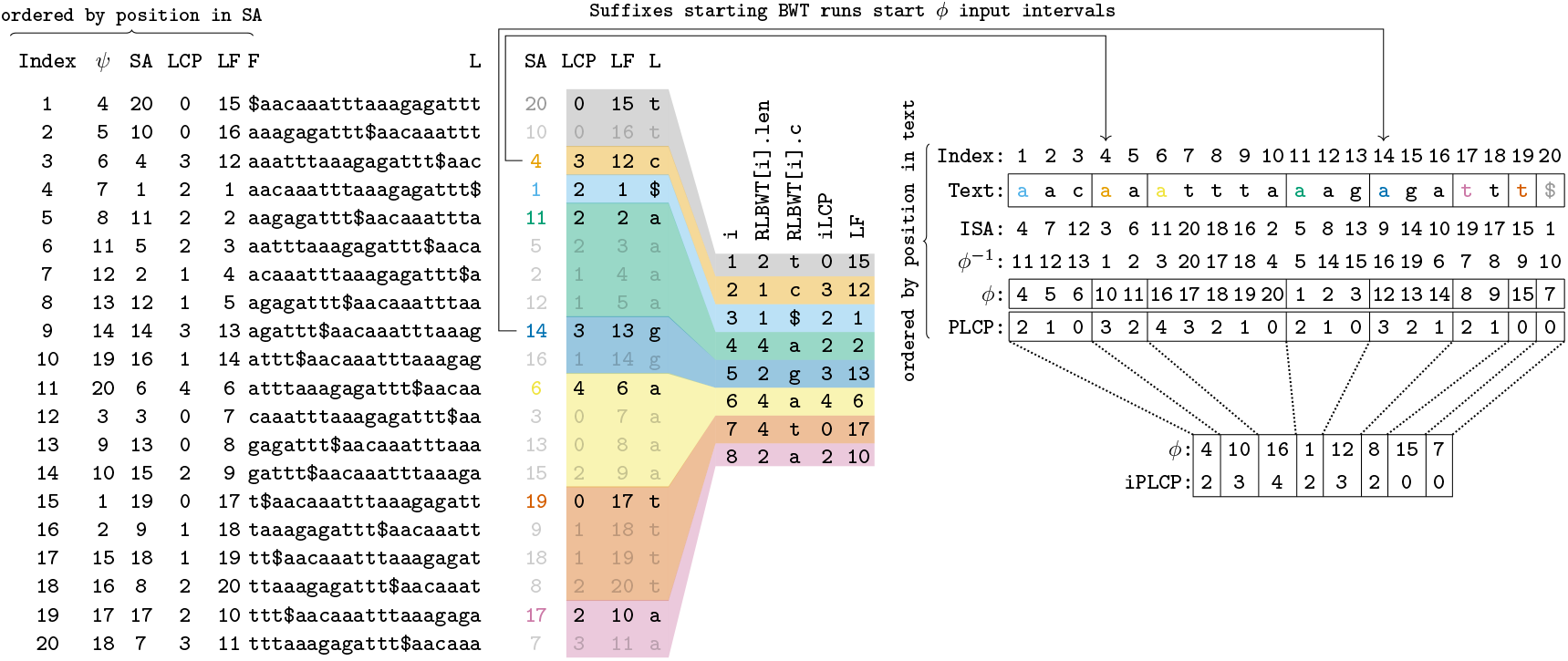
Relationships between uncompressed and run-length compressed *BWT* data structures

A run in the BWT of *T* is an interval [*i, j*] where *BWT* [*i*^′^] = *BWT* [*i*] for all *i*^′^ ∈ [*i, j*] and *BWT* [*I* − 1] ≠ *BWT* [*i*] and *BWT* [*j* + 1] ≠ *BWT* [*i*]. We use *r* to denote the number of runs in a BWT. Note *σ* ≤ *r* ≤ *n*. Then, the run length encoded Burrows-Wheeler transform, RLBWT, of *T* is an array of *r* 2-tuples (*c, len*) where the *k*-th run in the BWT, [*i*_*k*_, *j*_*k*_], is represented by the *k*-th value in the RLBWT as (*BWT* [*i*_*k*_], *j*_*k*_ − *i*_*k*_ + 1). *RLBWT* [*k*].*c* is the character of the *k*-th run and *RLBWT* [*k*].*len* is its length. For any run [*i, j*] in the BWT and *i*^′^ ∈ [*i, j*], *LF* [*i*^′^] = *LF* [*i*] + (*i*^′^ − *i*). It has also been shown that LF and *ϕ* are permutations with *r* intervals [Gagie et al., 2020, Nishimoto and Tabei, 2021]. Therefore, they can ∈ be computed with a move query in constant time and *O*(*r*) space [Nishimoto and Tabei, 2021]. It has also been shown that *PLCP* [*i*] can be computed in constant time within *O*(*r*) space during a move query on *ϕ* [Sanaullah et al., 2025]. A “run” in *ϕ* is an interval [*i, j*] where *ϕ*[*i*^′^] = *ϕ*[*i*] + (*i*^′^ − *i*) for all *i*^′^ ∈ [*i, j*]. Let *p*_*h*_ be the *h*-th smallest value| ∈ [1, *n*] s.t. *p*_*h*_ occurs in the SA at a position corresponding to the top of a run in the BWT. Recall the definition of run in the BWT, for *k* ∈ [1, *r*], [*i*_*k*_, *j*_*k*_] is the *k*-th run in the BWT. Then *h* = |{*k* |*k* ∈ [1, *r*] ∧ *SA*[*i*_*k*_] ≤ *p*_*h*_ and ∃*k* s.t. *p*_*h*_ = *SA*[*i*_*k*_]. I.E. *h* ∈ [1, *r*] and *p*_1_, *p*_2_, *p*_3_, …, *p*_*r*_ are the suffixes at the top of runs in the BWT sorted by starting position in the text (not sorted lexicographically). Then [*p*_*h*_, *p*_*h*+1_ − 1] is a “run” in *ϕ*. Furthermore, for any such run, *PLCP* [*i*^′^] = *PLCP* [*p*_*h*_] − (*i*^′^ − *p*_*h*_) for *i*^′^ ∈ [*p*_*h*_, *p*_*h*+1_ − 1] [Gagie et al., 2020, Sanaullah et al., 2025]. Finally, irreducible LCP (iLCP) values are those that occur at the top of runs in the BWT, and irreducible PLCP (iPLCP) values are the PLCP values of the suffixes that occur at the top of runs in the BWT. Figure 1 depicts run-length compressed data structures of a BWT.

#### 2.2.1 Coordinates

In the uncompressed BWT/LF (equivalently *ϕ, PLCP*), a “coordinate” is simply an integer *i*, and the associated data can be retrieved at position *i* in the BWT/LF array (respectively *ϕ, PLCP* array) in *O*(*n*) space and *O*(1) time. However, “coordinate” is not so simple in the run-length compressed BWT/LF (respectively *ϕ, PLCP*). Given *i*, it is not necessarily possible to retrieve *BWT* [*i*], *LF* [*i*] (respectively *ϕ*[*i*], *PLCP* [*i*]) in *O*(*r*) space and *O*(1) time. However, given *x*, the index of the input interval of LF (respectively, of *ϕ*) that contains position *i*, and *off*, the offset of position *i* within the input interval, it is possible to compute *LF* [*i*] (respectively, *ϕ*[*i*] and *PLCP* [*i*]) in *O*(*r*) space and *O*(1) time. Although *x* and *off* are sufficient to compute BWT/LF (equivalently, *ϕ, PLCP*), the original value *i* is often desired, such as when counting occurrences in a range or when reporting the location of a single occurrence. Therefore, we define a coordinate as a triple *crd* = (*i, x, off*). If the coordinate is a BWT/LF coordinate, it represents a position in the suffix array (*i*), and the triple is subject to the restriction *i*_*x*_ + *off* = *i* ≤ *i*_*x*+1_. If the coordinate is a *ϕ/PLCP* coordinate, it represents a position in the text, *i*, (or equivalently, a suffix, suffix *i*), and the triple is subject to the restriction *p*_*x*_ + *off* = *i* < *p*_*x*+1_. We refer to *i* by *crd*.*pos, x* by *crd*.*interval*, and *off* by *crd*.*offset*.

Although a coordinate allows the computation of *LF* (or *ϕ/PLCP*) once, it is frequently desired to compose the function calls (i.e. to compute *LF* [*LF* [*i*]], respectively, *ϕ*[*ϕ*[*i*]]). In order to do so, it is required to compute alongside *LF* [*i*] (or *ϕ*[*i*]), *x*^′^, where *x*^′^ is the index of the input interval of *LF* (or *ϕ*) that contains *LF* [*i*] (or *ϕ*[*i*]). As discussed in Section 2.1, the move data structure allows the maintenance of this value in *O*(*r*) space and *O*(1) time (also see relevant works: [Bertram et al., 2024, Brown et al., 2022, Nishimoto and Tabei, 2021]).

## 3 Methods

Although the traditional definition of a disjoint interval sequence is an ordered list of *r* pairs of integers in [1, *n*], (*p*_*i*_, *f* (*p*_*i*_)) where [*p*_*i*_, *p*_*i*+1_ − 1] is a conserved interval (define *p*_*r*+1_ = *n* + 1), we define disjoint interval permutation, a closely related definition that may be more useful in some cases. A *disjoint interval permutation* is a pair of ordered lists of integers, (*p, π*) where *π* is a permutation of [1, *r*] and *p* is a sorted list of *r* + 1 distinct integers in [1, *n* + 1] with *p*[*r* + 1] = *n* + 1. Call (*p, π*) the disjoint interval permutation that is equivalent to a disjoint interval sequence composed of pairs 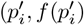). Then 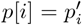 and *π*[*i*] = {|*p*[*j*] : *f* (*p*[*j*]) ≤ *f* (*p*[*i*])}|. *π* here is the inverse of *π* in the original move data structure paper [Nishimoto and Tabei, 2021]. For constant *n*, there is a clear one to one correspondence between disjoint interval permutations and disjoint interval sequences. A disjoint interval permutation can be easily converted to a disjoint interval sequence in *O*(*r*) space and time (see Algorithm 1 in the Supplementary Material). Furthermore, there are benefits to using disjoint interval permutations instead of disjoint interval sequences. For example, the disjoint interval permutation (and sequence) of *f*^−1^ can be easily computed from the disjoint interval permutation of *f* in *O*(*r*) space and time (see Algorithm 2 in the Supplementary Material). Therefore, the algorithms in this paper compute disjoint interval permutations. They can all be easily modified to compute disjoint interval sequences with unchanged time complexity since they all use Ω(*r*) space and time.

We describe algorithms that construct the disjoint interval permutations of various run-length Burrows-Wheeler transform (RLBWT) related functions given the RLBWT. In Section 3.1, we provide an algorithm for computing the disjoint interval permutations of *LF* and *ψ* in *O*(*r*) space and time given the RLBWT. In Section 3.2 we provide an algorithm for computing the interval permutation of *ϕ* and *ϕ*^−1^ in *O*(*r*) space and *O*(*n*) time given the RLBWT and the move data structure of *ψ*. Finally, in Section 3.3, we provide an algorithm for computing all irreducible PLCP values in *O*(*r*) space and *O*(*n*) time given the RLBWT and the move data structure of *ψ*.

### 3.1 *LF* and *ψ* Construction

Here, we present algorithms for the construction of the disjoint interval permutations of *LF* and *ψ* of a text in *O*(*r*) space and time given its RLBWT. As a result, we obtain *O*(*r*) space and time algorithms for computing the disjoint interval sequences of *LF* and *ψ* given the RLBWT. Finally, providing the disjoint interval sequence to the move data structure construction algorithm described by Bertram et al. [2024] results in *O*(*r*) space *O*(*r* log *r*) time algorithms for *LF* and *ψ* move data structure construction given the RLBWT.

We first describe the algorithm for constructing the disjoint interval permutation of *LF* given the RLBWT in *O*(*r*) time and space. The algorithm takes two linear sweeps of the RLBWT. In the first sweep, the number of runs of each character in the RLBWT is accumulated. After this sweep, an exclusive prefix sum in lexicographical order is done to obtain *C*_*r*_[*c*] for all *c* ∈ Σ, where *C*_*r*_[*c*] is total number of runs of characters *c*^′^ < *c* in the BWT. Finally a second linear sweep of the RLBWT is done, where the number of runs seen so far is maintained for each character in Σ, (*rank*(*RLBWT, c, i*)). The start position of each run is also maintained (*p*^′^). Then, *p*[*i*] = *p*^′^ and *π*[*i*] = *C*_*r*_[*RLBWT* [*i*].*c*] + *rank*(*RLBWT, RLBWT* [*i*].*c, i*) can be set in constant time per *i* during the scan. See Algorithm 3 in the Supplementary Material for pseudocode of this algorithm. A depiction of this algorithm is available in Figure 2.

**Figure 2.**
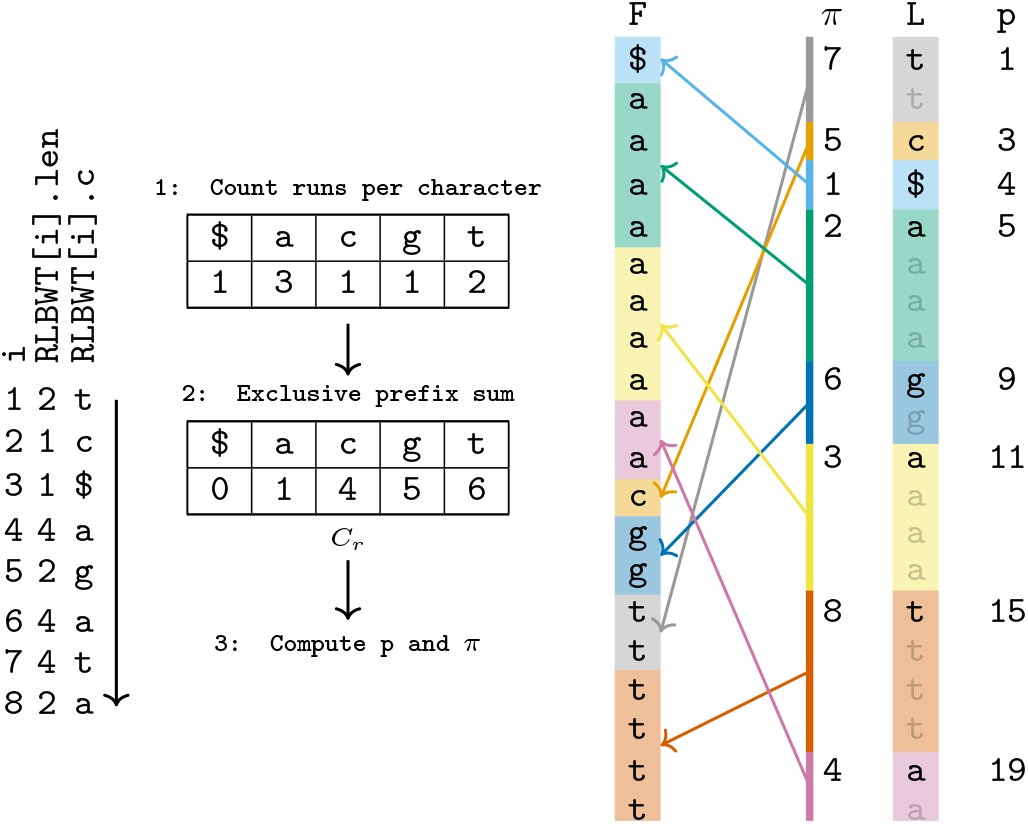
Construction of LF disjoint interval permutation in *O*(*r*) space given an RLBWT. *T* = *aacaaatttaaagagattt*$. The expanded view on the right is for illustrative purposes, it is not present in memory. *L* represents the BWT of *T, L* = *BWT* (*T*).

To obtain the *ψ* disjoint interval permutation, the *LF* disjoint interval permutation can be inverted in *O*(*r*) space and time with Algorithm 2 in the Supplementary Material. To obtain the *ψ* or *LF* disjoint interval sequences in *O*(*r*) space and time, the corresponding disjoint interval permutation can be converted in *O*(*r*) space and time with Algorithm 1 in the Supplementary Material. Finally, to construct the move data structures of *LF* and *ψ* in *O*(*r* log *r*) time and *O*(*r*) space, the corresponding interval sequence can be provided as input to the construction algorithm of Bertram et al. [2024].

### 3.2 ϕ and *ϕ*^−1^ Construction

Here, we present algorithms for the construction of the disjoint interval permutations and sequences of *ϕ* and *ϕ*^−1^ in *O*(*n*) time and *O*(*r*) space given the RLBWT and a move data structure for *ψ*. This also implies *O*(*n* + *r* log *r*) time and *O*(*r*) time algorithms for computing the *ϕ* and *ϕ*^−1^ move data structures given the RLBWT or text.

We first describe the algorithm for constructing the *ϕ* disjoint interval permutation, (*p, π*), given the RLBWT and a move data structure for *ψ*. First, the disjoint interval permutation of *ψ* is obtained in *O*(*r*) time and space using the algorithm described in the previous section. The algorithm then proceeds in three steps. First, the RLBWT is converted into RF^′^ in *O*(*r*) space and time, where RF and RF^′^ are F split into runs according to input intervals of *ψ* disjoint interval sequence and move data structure respectively. Second, an *O*(*n*) time *O*(*r*) space traversal of the BWT is completed to compute *p* and *SA*_*inp*_. Finally another *O*(*n*) time *O*(*r*) space traversal of the BWT is completed to compute *π*.

The first step of the algorithm, converting the RLBWT to RF^′^ is discussed in Section S1.1 in the Supplementary Material. The second step of the algorithm is an *O*(*n*) time and *O*(*r*) space traversal of the BWT in text order that computes *p* and *SA*_*inp*_. *p* is the first component of (*p, π*), the disjoint interval permutation of *ϕ. SA*_*inp*_ is an array of size *O*(*r*). Call *s* the suffix of *T* that occurs at the top of *RF* [*i*] for some *i* ∈ [1, *r*], then there exists *i*^′^ s.t. *s* occurs at the top of RF^′^[*i*^′^]. Then *s* is the end of a *ϕ* input interval, since *s* + 1 is the start of a *ϕ* input interval. Then define *SA*_*inp*_[*i*^′^] as the index of the input interval of *ϕ* that *s* occurs at the end of. This step begins with obtaining *crd*_*ψ*_, the *ψ* coordinate corresponding to suffix *n* of the text. Since each terminal character $_∗_ is contained in its own input interval of size 1 of the *ψ* move data structure, (*N, N*, 1) is the coordinate of suffix *n* in the *ψ* move data structure, where *N* is the number of terminal characters in *T*. Then, *n ψ* operations are computed by *crd*_*ψ*_ = ℳ_*ψ*_(*crd*_*ψ*_) while maintaining the current suffix (*s*) and number of times previous suffixes have been at the top of runs in RF (*tops*). Then when *crd*_*ψ*_ is at the top of a run in the RF, *p* and *SA*_*inp*_ are updated appropriately.

The third and final step of the algorithm is another *O*(*n*) time and *O*(*r*) space traversal of the BWT in text order that computes *π*, the second component of the disjoint interval permutation of *ϕ*. This is done by again starting from *crd*_*ψ*_ = (*N, N*, 1) and computing *n ψ* operations. This time, the number of times previous suffixes have been at the bottom of a run in the BWT is maintained (*bots*). When *crd*_*ψ*_ is at the bottom of a run in the BWT, a *π* value can be computed using *SA*_*inp*_. The pseudocode for this algorithm can be seen in Algorithm 6 in the Supplementary Material. Figure 3 depicts part of the first *O*(*n*) time traversal of this algorithm.

**Figure 3.**
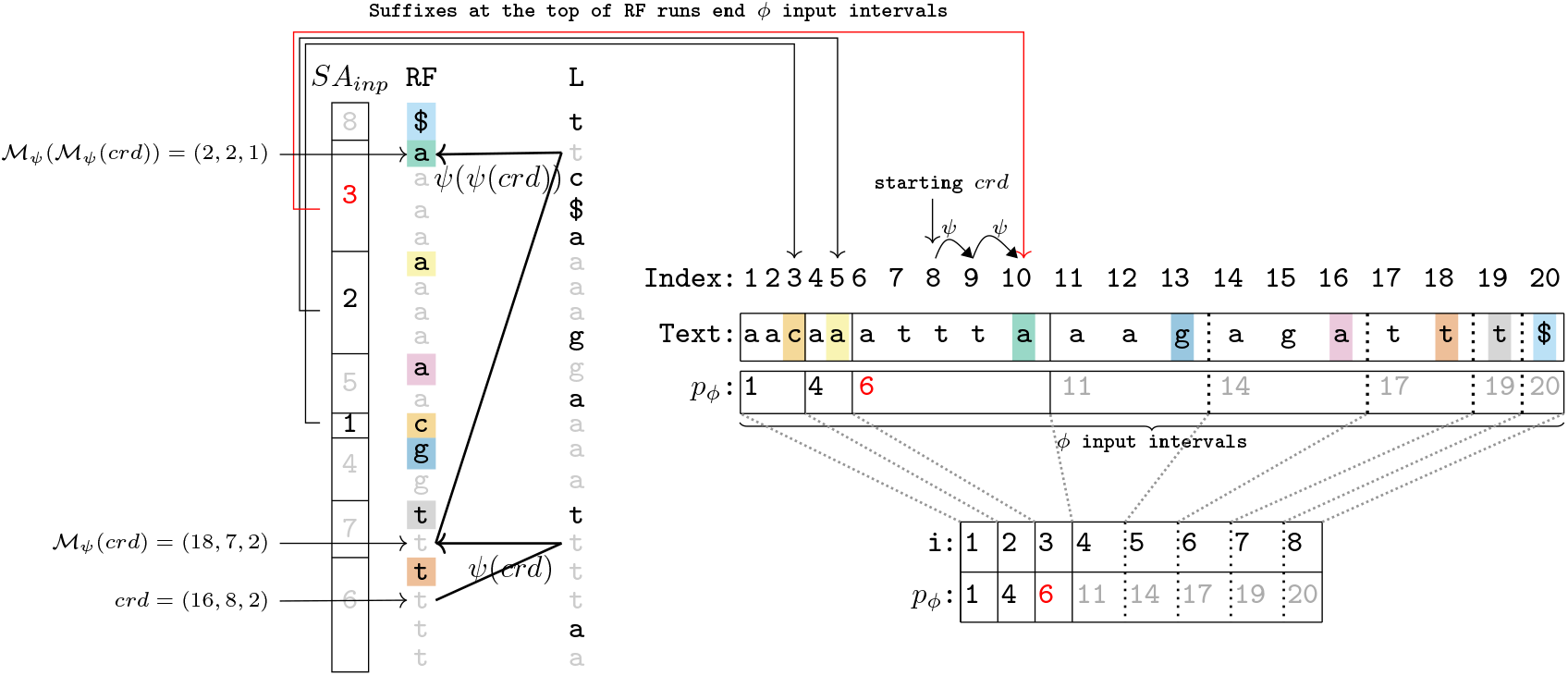
*ϕ* construction, computing *p* and *SA*_*inp*_. Black, red, and gray values were computed in the past, present, and future respectively.

**Figure 4.**
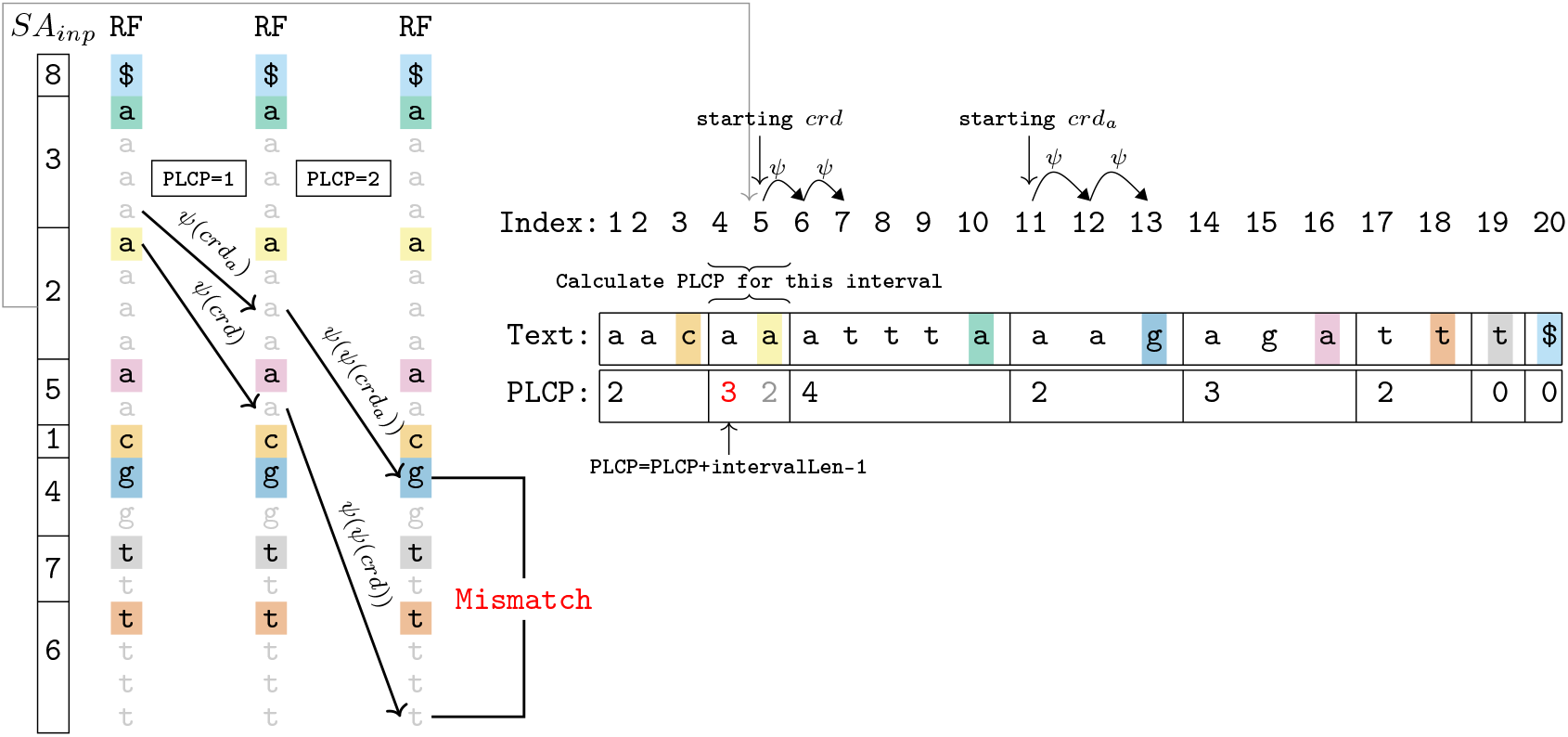
Computing irreducible PLCP algorithms with the naive algorithm

The *ϕ*^−1^ disjoint interval permutation can be obtained from the *ϕ* disjoint interval permutation with an additional *O*(*r*) time and space using Algorithm 2 in the Supplementary Material. This results in an overall *O*(*n*) time *O*(*r*) space algorithm for obtaining the *ϕ*^−1^ disjoint interval permutation given the RLBWT and a move data structure for *ψ*. These result in algorithms for obtaining the disjoint interval sequences of *ϕ* and *ϕ*^−1^ with the same time and space complexities given the same inputs by applying Algorithm 1 in Supplementary Material. Lastly, the move data structures of *ϕ* and *ϕ*^−1^ can be obtained in *O*(*n* + *r* log *r*) time and *O*(*r*) space given only RLBWT by first generating the *ψ* move data structure with the algorithm from Section 3.1, constructing the disjoint interval sequence by the algorithm described in this section, and then providing the disjoint interval sequences to the construction algorithm of Bertram et al. [2024]. Note that the *r* log *r* term in this time complexity is only from balancing the disjoint interval sequences.

### 3.3 Irreducible PLCP and LCP Computation

Here, we provide algorithms for computing irreducible PLCP and LCP values of a text in run length compressed space given the RLBWT. First, we provide a naive algorithm that achieves *O*(*n* log *δ*) ∈ *O*(*n* log *r*) ∈ *O*(*n* log *n*) time and *O*(*r*) space given the RLBWT and a *ψ* move data structure. Then we propose an algorithm that achieves linear time through a sampling of the ISA. This algorithm computes irreducible PLCP and LCP values of the text in *O*(*n*) time and *O*(*r*) space given the RLBWT and a *ψ* move data structure. These result in *O*(*n* log *r*) ∈ *O*(*n* log *n*) and *O*(*n* + *r* log *r*) time algorithms respectively for obtaining the irreducible PLCP or LCP values in *O*(*r*) space given the RLBWT (or given just the text [Nishimoto et al., 2022]). In the description below, we only compute PLCP values, but these can be converted to LCP values at the top of runs in *O*(*r*) time and space given the RLBWT and *SA*_*inp*_, see Section S1.2 in the Supplementary Material.

#### 3.3.1 Naive Algorithm

The naive algorithm is a variation of the brute force approach of Kärkkäinen et al. [2009]. Our algorithm first obtains the disjoint interval permutation of *ϕ* and *SA*_*inp*_ in *O*(*r*) time and space as described in Section 3.2. Then, the algorithm starts at the top of every run in RF. For every run *i* of RF, it does the following: it maintains two *ψ* coordinates: *crd* and *crd*_*a*_. Initially, *crd* is set to the top of the current run and *crd*_*a*_ is set to the position just above *crd*. Then, *crd* = ℳ_*ψ*_(*crd*) and *crd*_*a*_ = ℳ_*ψ*_(*crd*_*a*_) is computed as long as *RF*^′^[*crd*.*interval*] = *RF*^′^[*crd*_*a*_.*interval*]. The number of times *RF*^′^[*crd*.*interval*] = *RF*^′^[*crd*_*a*_.*interval*] plus the length of the *SA*_*inp*_[*i*]-th input interval of *ϕ* minus one is the PLCP value of the *SA*_*inp*_[*i*]-th input interval of *ϕ*. See Figure 3 for a depiction of one the *iPLCP* value computations of this algorithm. For pseudocode of this algorithm, see Algorithm 7 in the Supplementary Material. The number of *ψ* move operations completed in total for all runs is at most two times the sum of the irreducible PLCP values. The irreducible PLCP values are a permutation of the irreducible LCP values, and the sum of the irreducible LCP values has been shown to be *O*(*n* log *δ*) ∈ *O*(*n* log *r*) ∈ *O*(*n* log *n*) [Kärkkäinen et al., 2015, Kempa and Kociumaka, 2022]. Using the algorithms from Section 3.2, computing the *ϕ* disjoint interval permutation and *SA*_*inp*_ takes for *O*(*n*) time and *O*(*r*) space given the RLBWT and a *ψ* move data structure. Therefore, these result in algorithms for computing the irreducible PLCP and LCP values in *O*(*n* log *δ*) ∈ *O*(*n* log *r*) ∈ *O*(*n* log *n*) time and *O*(*r*) space given the RLBWT and a *ψ* move data structure, or *O*(*n* log *δ* + *r* log *r*) ∈ *O*(*n* log *r*) ∈ *O*(*n* log *n*) time and *O*(*r*) space given the text.

#### 3.3.2 Linear Time Algorithm

Here, we propose an algorithm for *O*(*r*) space and *O*(*n*) time computation of all irreducible PLCP values given the RLBWT and a move data structure for *ψ*. This algorithm is a variation of a classic LCP computation algorithm by Kasai et al. [2001] and its extension by Kärkkäinen et al. [2009] which uses *ϕ* and PLCP values. The key idea behind our algorithm is an *O*(*r*) sampling of the ISA. First *SA*_*inp*_ and the disjoint interval permutation of *ϕ* are computed in *O*(*n*) time and *O*(*r*) space with the algorithm described in Section 3.2. Then *SA*_*inp*_ is inverted to compute 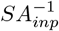 in *O*(*r*) time and space. Then, an *O*(*n*) time *O*(*r*) space traversal is done to compute *r* equidistant ISA samples (i.e 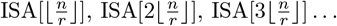). The values stored for these ISA samples are the *ψ* coordinates where the SA values occur. Then, a traversal through the input intervals of the *ϕ* in text order is completed. During this traversal, a *ψ* coordinate of the smallest unmatched suffix is maintained, *crd*. Initially, *crd* is the *ψ* coordinate where suffix 1 of the text occurs. We also maintain *s*, the suffix occurring at *crd*. Initially, *s* = 1.

Having already computed the *iPLCP* values from 1 to *k*, we compute *iPLCP* [*k*+1] as follows. *iPLCP* [*k*+ 1] is the matching length between suffixes *p*_*ϕ*_[*k* + 1] and *ϕ*(*p*_*ϕ*_[*k* + 1]). It is also known that *p*_*ϕ*_[*k* + 1] + *iPLCP* [*k* + 1] ≥ *p*_*ϕ*_[*k*] + *iPLCP* [*k*] (Lemma 3 in Kasai et al. [2001]). Therefore, we first obtain *crd*_*a*_, the *ψ* coordinate corresponding to the position where suffix *ϕ*(*p*_*ϕ*_[*k* + 1]) + *s* − *p*_*ϕ*_[*k* + 1] is located in the BWT (aka ISA[*ϕ*(*p*_*ϕ*_[*k* + 1]) + *s* − *p*_*ϕ*_[*k* + 1]]). This coordinate is recovered in 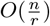 time by starting from the closest preceding ISA sample and computing at most 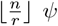 operations. Finally, as long as *RF*^′^[*crd*.*interval*] = *RF*^′^[*crd*_*a*_.*interval*], *crd* is set to ℳ_*ψ*_(*crd*), *crd*_*a*_ is set to ℳ_*ψ*_(*crd*_*a*_) and *s* = *s* + 1. Then, *iPLCP* [*k* + 1] = *s* − *p*_*ϕ*_[*k* + 1].

In this algorithm, throughout the whole sweep from *k* = 1 → *r*, at most *n* total *ψ* operations are computed on *crd* because *s* is an integer that goes from 1 to n and is incremented exactly once for every *ψ* operation computed on *crd*. Therefore, at most *n ψ* operations are computed on *crd*_*a*_ since a *ψ* operation is computed on *crd*_*a*_ iff a *ψ* operation is computed on *crd*. Finally, *r crd*_*a*_ values are initialized, each with at most 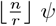 operations needed to initialize it from the ISA samples, resulting in a total *O*(*n*) *ψ* operations. Therefore, this algorithm computes all irreducible PLCP values (or irreducible LCP values) in *O*(*r*) space and *O*(*n*) time given the RLBWT and a move data structure for *ψ*. Finally, this implies an *O*(*n* + *r* log *r*) time and *O*(*r*) space algorithm for computing all irreducible PLCP and LCP values given the text (with the RLBWT construction algorithm of Nishimoto et al. [2022] and move data structure construction algorithm of Bertram et al. [2024]). This allows optimal time computation of the LCP array for repetitive 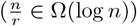) datasets while using *O*(*r*) additional space; this is the first such algorithm to achieve these bounds.

## 4 Results

We demonstrate that the algorithms presented in this work substantially decrease resource usage for computation of RLBWT support structure relative to state of the art algorithms in practice. In order to do so, we benchmark LCP summary computation on DNA datasets ranging from a few billion basepairs to a few trillion basepairs. The implementation of the algorithms described in this paper is available as TeraLCP, located within TeraTools. TeraLCP is parallelized by computing each algorithm per string in the text in parallel. See the parallelization benchmark in Supplementary Figure S1 and Table S5.

In benchmarks comparing in-memory LCP summary computation in compressed space, there only two competitive tools the authors are aware of: pfp-thresholds [Rossi et al., 2022] and rlbwt2lcp [Prezza and Rosone, 2021].

Five datasets are utilized for the experiments: mtb152, chr19.1000, human100, human472, and CommonBacteria. mtb152 is 152 M. tuberculosis genomes. chr19.1000 is 1,000 chromosome 19 haplotypes from 1000G Phase 3 [The 1000 Genomes Project Consortium, 2015], generated by bcftools consensus. human100 and human472 are collections of 100 and 472 human haplotype assemblies respectively. The assemblies from HPRC release 1 are contained in human100 and the assemblies from HPRC release 2 are contained in human472 [Liao et al., 2023]. CommonBacteria is a collection of over a million of bacterial genome assemblies, obtained from AllTheBacteria v0.2 [Hunt et al., 2025].Since all datasets are DNA, the text that the RLBWT is constructed on contains the original string and its reverse complement for every string in the dataset. Statistics and sources for these datasets can be seen in Supplementary Table S4. See Section S1.3 in the Supplementary Material for computing environment details for all experiments.

We present three experiments to evaluate the performance of TeraLCP. We focus on LCP summaries of the text since they are the most expensive RLBWT support structures to compute in practice and have the least developed algorithms and tools. We first show that TeraLCP, significantly decreases running time and peak memory relative to pfp-thresholds and rlbwt2lcp. Next, we show TeraLCP enables the computation of LCP summaries of terabase scale datasets in a typical university compute node. Finally, we show that the memory usage of TeraLCP increases slower than that of pfp-thresholds and rlbwt2lcp as repetitive datasets get larger.

### 4.1 Text to Minimum LCP per Run

In this experiment, we benchmark computing the minimum LCP value per run in the BWT. This is the only LCP summary that rlbwt2lcp, TeraLCP, and pfp-thresholds all share in common. This is done for the mtb152 and chr19.1000 datasets. The results can be seen in summary form in Table 2 and in more detail in Supplementary Table S7. The results of rlbwt2lcp and TeraLCP include RLBWT construction time and peak memory using lyndongrammar [Olbrich, 2025] with 16 threads. RLBWT construction benchmarking (available in Supplementary Table S2) was conducted to select the RLBWT construction method. lyndongrammar was selected because it provided a good balance of low memory usage and fast run time out of the tested tools. TeraLCP significantly improves on both running time and peak memory on both rlbwt2lcp and pfp-thresholds in both tested datasets. It uses 4.5 times and 5.8 times less memory on mtb152 and 14.6 times and 27.2 times less memory on chr19.1000 than pfp-thresholds and rlbwt2lcp respectively. TeraLCP also takes 7.5 times and 20.2 times less time on mtb152 and 5.4 times and 29.1 times less time on chr19.1000 than pfp-thresholds and rlbwt2lcp respectively.

**Table 2:**
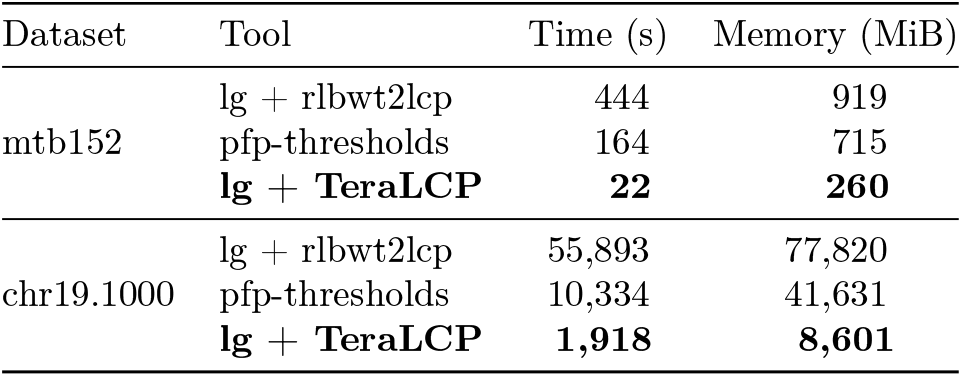
Minimum LCP per run computation.

| Dataset | Tool | Time (s) | Memory (MiB) |
| --- | --- | --- | --- |
| mtb152 | lg + rlbwt2lcp | 444 | 919 |
|  | pfp-thresholds | 164 | 715 |
|  | **lg + TeraLCP** | **22** | **260** |
| chr19.1000 | lg + rlbwt2lcp | 55,893 | 77,820 |
|  | pfp-thresholds | 10,334 | 41,631 |
|  | **lg + TeraLCP** | **1,918** | **8,601** |

### 4.2 LCP Summaries of Terabase Scale Datasets

We first present the memory usage of the three LCP summary generating tools (pfp-thresholds, rlbwt2lcp, and TeraLCP) on all five datasets (mtb152, chr19.1000, human100, human472, and CommonBacteria). RLBWT generation (with sequential grlbwt [Díaz-Domínguez and Navarro, 2023]) used less memory than all LCP summary generating tools on all datasets. Due to compute limitations, programs run on larger datasets were run on different environments. Therefore, the run time results are not directly comparable and have been left out. The results can be seen in Table 3. RLBWT generation with sequential grlbwt [Díaz-Domínguez and Navarro, 2023] had a lower memory consumption than all LCP summary tools for mtb152 and chr19.1000. For human100 and CommonBacteria, rlbwt generation with ropebwt3 takes ≤ 83 GiB according to Li [2024]. For human472, rlbwt generation with ropebwt3 likely takes *<* 170 GiB according to the 99 GiB memory usage on human320 in [Li, 2024]. The memory usage of rlbwt2lcp is estimated using the formula from its github for human100, human472, and CommonBacteria. The same formula approximates the actual memory usage well on mtb152 and chr19.1000: it estimates 0.92 GiB and 79.14 GiB respectively. pfp-thresholds was not run on CommonBacteria due to the authors only having access to machines with < 3 TiB RAM. TeraLCP uses significantly less memory than pfp-thresholds and rlbwt2lcp on all tested datasets. This enables the computation of LCP summaries of large biologically relevant datasets (e.g. human472 and CommonBacteria) on typical university compute nodes.

**Table 3:**
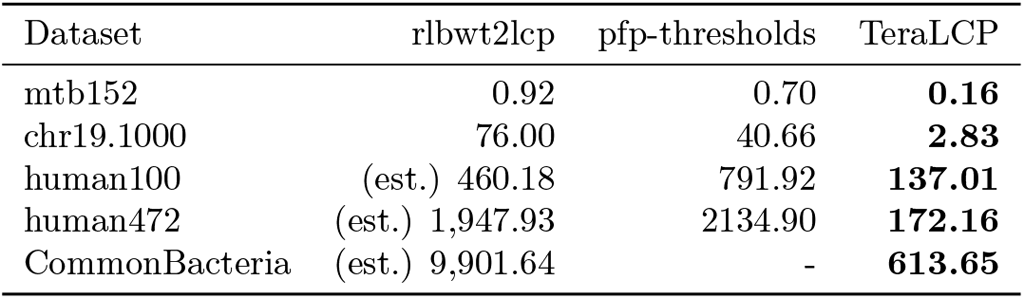
Peak Memory Usage (GiB) for LCP Summary Tools.

| Dataset | rlbwt2lcp | pfp-thresholds | TeraLCP |
| --- | --- | --- | --- |
| mtb152 | 0.92 | 0.70 | **0.16** |
| chr19.1000 | 76.00 | 40.66 | **2.83** |
| human100 | (est.) 460.18 | 791.92 | **137.01** |
| human472 | (est.) 1,947.93 | 2134.90 | **172.16** |
| CommonBacteria | (est.) 9,901.64 | - | **613.65** |

### 4.3 Memory Usage Scaling as Datasets Grow

Finally, we compare the memory usage of TeraLCP, rlbwt2lcp, and pfp-thresholds as the size of the text grows. For the mtb152, chr19.1000, and human100 datasets the following experiment was conducted. Call the number of strings in the original dataset *N*, then for *x* ∈ 1, 2, 4, 8, 16, 32, 64, a random 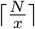 of the strings are selected. Each random selection of strings is a subset of larger random selections of strings for the same dataset. For each *x*, the RLBWT is constructed, then TeraLCP, rlbwt2lcp, and pfp-thresholds are run on the RLBWT, BWT, and the text respectively. rlbwt2lcp was not run on human100 subsets due to it being slow, however its memory usage is estimated by the formula on its github. The results for chr19.1000 and human100 are plotted in Figure 5 and mtb152 is plotted in Supplementary Figure S2. For all datasets, RLBWT construction (with single threaded grlbwt) used less memory than rlbwt2lcp and TeraLCP. On mtb152, TeraLCP uses less memory on the full dataset than pfp-thresholds uses on 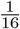 of the data. On chr19.1000 and human100, TeraLCP uses less memory on the full dataset than pfp-thresholds does on 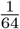 of the data. Finally, the memory usage of TeraLCP grows much slower than that of pfp-thresholds and rlbwt2lcp. From 50% of the data to 100% of the data, the memory usage of rlbwt2lcp increases by 81%, 97%, and 79% on mtb152, chr19.1000, and human100 respectively. For pfp-thresholds, the same values are 59%, 80%, and 43% respectively. However, for TeraLCP, these increases are 4%, 13%, and 7% respectively. The full data is available in Supplementary Table S6.

**Figure 5.**
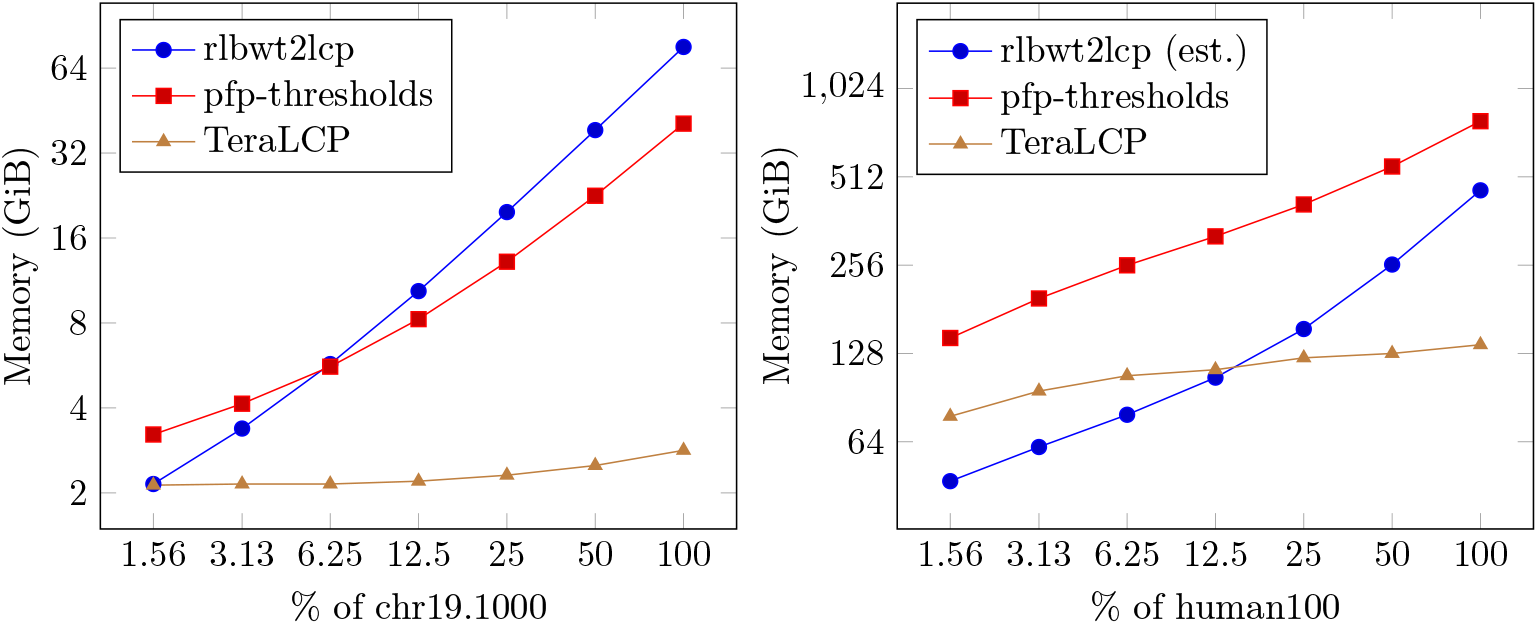
Memory scaling of TeraLCP, rlbwt2lcp, and pfp-thresholds

## 5 Discussion

We have presented *O*(*r*) space and *O*(*n* + *r* log *r*) time algorithms for computing many RLBWT support structures including LF, *ψ, ϕ, ϕ*^−1^, and irreducible LCP and PLCP values given the RLBWT. In conjunction with the algorithm of Nishimoto et al. [2022], this results in algorithms computing the same outputs directly from the text with the same time and space complexities. Note that the majority of datasets that these structures are computed on are very repetitive, with 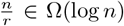. This results in linear *O*(*n*) time and run-length space *O*(*r*) for those repetitive datasets. Also note that given these support structures, many other important summaries can be computed straightforwardly in *O*(*r*) space and *O*(*n*) time. Our algorithms trivially extend to computing minimum LCPs per run and thresholds used in matching statistic computation [Rossi et al., 2022]. We provide an implementation of the algorithms discussed here as TeraLCP, contained in the TeraTools repository. Then, we show that the memory usage of TeraLCP scales much better in practice than previous LCP summary computing methods. It is also much faster than previous methods given multiple cores. In all datasets we tested, there is an RLBWT construction tool s.t. using that tool in conjunction with TeraLCP results in both lower run time and memory usage than both pfp-thresholds and rlbwt2lcp.

Barring RLBWT construction, the *r* log *r* term in the time complexities of the algorithms presented in this paper is only from one component of the algorithm: balancing the disjoint interval sequences. The authors believe an *O*(*r*) time and space algorithm for balancing disjoint interval sequences exists given the disjoint interval permutations (which the algorithms described here produce). Such an algorithm would improve the time complexities of the algorithms presented here to *O*(*n*) time and *O*(*r*) space in general. We leave the description of this algorithm as future work. Furthermore, the performance of TeraTools still has room to improve. Currently TeraTools does not employ latency hiding techniques that have proven fruitful in other indexes [Zakeri et al., 2024]. Removing the cache miss bottleneck may significantly speed up the tool. Its current parallelization method is naive and may be improved. Furthermore, its memory consumption may be reduced by utilizing sparse bit vectors or capping maximum interval length [Brown et al., 2022]. Before this work, RLBWT support structure construction (particularly LCP related) was a significant bottleneck for lossless compressed full text index construction and all downstream analyses. After this work, RLBWT support structure construction is no longer the bottleneck, all RLBWT construction tools use more memory or much more time than TeraLCP for large repetitive datasets.

## 6 Acknowledgments

This work was supported by the National Institutes of Health under grants R01HG010086 and R01HG011392.

## Notes

https://github.com/ucfcbb/TeraTools

### Competing Interest Statement

The authors have declared no competing interest.

