## Supplementary Material for "RLBWT-Based LCP Computation in Compressed Space for Terabase-Scale Pangenome Analysis"

### S1 Supplementary Text

#### S1.1 RF and RF′

Let $(p_\psi, \pi_\psi)$ be the disjoint interval permutation of $\psi$. RF is the run-length encoded F array, according to where runs in the BWT map into the F array. Therefore, RF is an array of $r$ pairs $(c, len)$ where $RF[i] = RLBWT[\pi_\psi[i]]$. RF′ is the RF split into intervals corresponding to the input intervals of the move data structure of $\psi$. Therefore RF′ is an array of pairs $(c, len)$ of size $O(r)$ (more precisely, $\leq 2r$ pairs), and each interval of RF′ is a run of characters of the same value (possibly a subrun of a run in RF). Therefore, for every run RF[$i$] in F, there is a corresponding range of intervals RF′$[i', j']$ in RF′ that represent it ($i'$ may equal $j'$); then RF$[i].len = \sum_{k'=i'}^{j'}$RF′$[k'].len$ and $\forall k' \in [i', j']$, RF$[i].c =$RF′$[k'].c$. Finally, the runs occur in the same order in RF and RF′; for runs RF$[i_1]$ and RF$[i_2]$ represented by RF′$[i'_1, j'_1]$ and RF′$[i'_2, j'_2]$ respectively, $i_1 < i_2 \iff i'_1 < i'_2$. Then RF′ is computed from RF and $\pi_\psi$ in $O(r)$ space and time by first splitting $\pi_\psi$ according to the input intervals of the psi move data structure, generating $\pi'_\psi$. Finally RF′ is generated by a linear scan through $\pi'_\psi$, using the lengths of the corresponding input intervals in the move data structure.

The move data structure as described in [1] is constructed on a balanced interval sequence and is composed of two components: $D_{pair}$ and $D_{index}$. $D_{pair}$ is an array of length $r$ that contains the representation of the balanced interval sequence: $D_{pair}[i] = (p_i, q_i)$. Given the balanced interval sequence (obtained from its move data structure) and the (unbalanced) disjoint interval permutation of $\psi$, the balanced disjoint interval permutation of $\psi$ is obtained from the method described above. Pseudocode of the process can also be seen in Algorithm 4.

#### S1.2 Irreducible PLCP to Irreducible LCP conversion

Here we describe how to convert irreducible PLCP values to irreducible LCP values given $SA_{inp}$. In Section 3.3 of the main paper, we provide algorithms for computing the irreducible PLCP. I.E. for each input interval of the disjoint interval permutation of $\phi$, the PLCP value at the start of input interval. Since the suffixes at the start of the input intervals of $\phi$ correspond are exactly the suffixes at the tops of runs in the RLBWT, all that is needed to convert irreducible PLCP to irreducible LCP is a mapping (call it $SA_{toRun}$) from input intervals of $\phi$ to the runs in the RLBWT s.t. the suffix at the top of run $SA_{toRun}[i]$ of the BWT and the suffix at the start of input interval $i$ of the disjoint interval permutation of $\phi$ are the same suffix. $SA_{inp}$ already maps $\psi$ input intervals to $\phi$ input intervals, however it maps the suffixes at the ends of $\phi$ input intervals to the $\phi$ input intervals that contain them. Furthermore, we desire the inverse mapping, so we invert $SA_{inp}$ to obtain $SA^{-1}_{inp}$, the mapping from $\phi$ input intervals to the $\psi$ input intervals s.t. the suffix at the end of $\phi$ input interval $k-1$ is at the top of $\psi$ input interval $SA^{-1}_{inp}[k-1]$. We can then use $\pi_\psi$, the permutation of the disjoint interval permutation of $\psi$. If the suffix at the top of input interval $j$ of $\psi$ ends the $\phi$ input interval $t$, then the suffix at the top of input interval $\pi_\psi[j]$ of LF (i.e. the $j$-th run of the BWT) begins the $t+1$-th input interval of $\phi$. Therefore, we obtain $SA_{toRun}$ as $SA_{toRun}[(k \mod r) + 1] = \pi_\psi[SA^{-1}_{inp}[k]]$. Finally the irreducible LCP value at the top of run $i = SA_{toRun}[k]$, $iLCP[i]$, can be computed by $iLCP[i] = iPLCP[k]$. $SA^{-1}_{inp}$ can be inverted from $SA_{inp}$ in $O(r)$ time and space and $\pi_\psi$ can be computed from the RLBWT in $O(r)$ time and space. Therefore $SA_{toRun}$ can be obtained in $O(r)$ time and space given $SA_{inp}$ and $\pi_\psi$. Then, $SA_{toRun}$ can be obtained in $O(r)$ time and space. Therefore, $iLCP$ can be obtained in $O(r)$ time and space given $iPLCP$ and $SA_{inp}$ and the RLBWT.

#### S1.3 Environments

See Table S3 for details for the computing environments. The only program run on aws was pfp-thresholds on human472. The RLBWT construction benchmarks in Table S2 were run on c0-4. Memory scaling benchmarks in Figure 5 and Table S6 were run on ec326 for mtb152 and chr19.1000 and were run on c0-5 for human100. The parallelization benchmark in Figure S1 and Table S5 was run on c0-5. The minimum LCP benchmark of Table S7 was run on c0-5. The mtb152 and chr19.1000 runs for all LCP summary tools in Table 3 were run on ec326. The TeraLCP results in Table 3 for human100, human472, and CommonBacteria were run on c0-5. The pfp-thresholds result for human472 in Table 3 was run on aws.

#### S2 Supplementary Figures

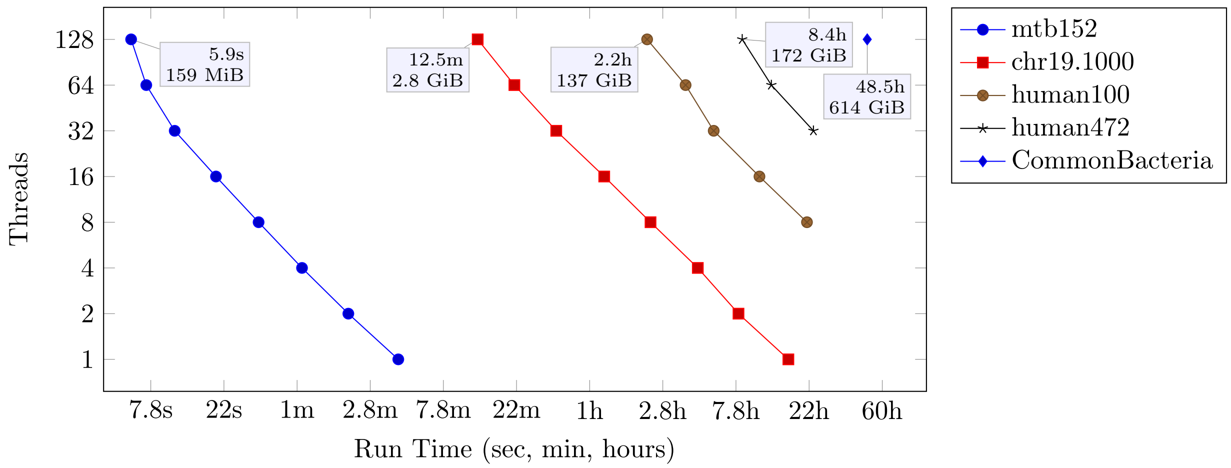


Figure S1: Run time of TeraLCP for various datasets and threads. The peak memory usage for each dataset was almost constant with respect to the number of threads used ($\frac{\max - \min}{\min} < 0.01$ for all datasets). See the equivalent precise numerical data in Table S5.

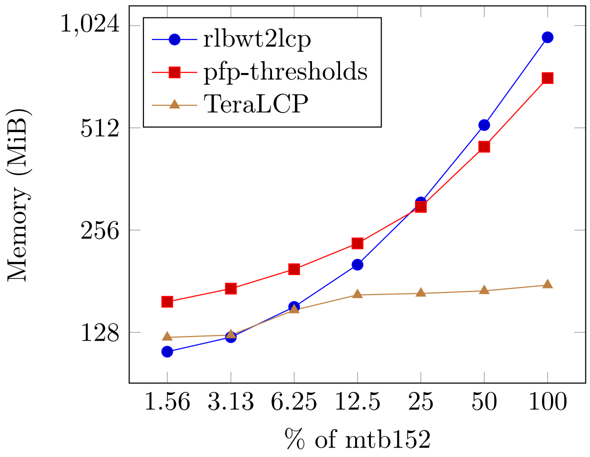


Figure S2: Peak memory usage of rlbwt2lcp, pfp-thresholds, and TeraLCP on subsets of mtb152

### S3 Supplementary Tables

Table S1: RLBWT construction algorithms

| Algorithm | Input | Output | Time Complexity | Space Complexity |
|---|---|---|---|---|
| grlbwt [2] | Text | RLBWT | $O(n + N \log n_{max})^{\dagger}$ | $O(n + N \log n_{max})$ |
| CMS-BWT [3] | Text$^{\ddagger}$ | RLBWT | $O(n \log \kappa + n \log \lvert\mathcal{R}\rvert + \lvert\mathcal{R}\rvert)$ | $O(\kappa + \lvert\mathcal{R}\rvert)$ |
| ropebwt3 [4] | Text | RLBWT | $O(n \log r)$ | $O(r)$ |
| r-comp [5] | Text | RLBWT | $O(n)^{\S}$ | $O(r)$ |
| lyndongrammar [6] | Text | LGram | $O(n \log g + g \log^2 g)$ | $O(g)$ |
| lyndongrammar [6] | Text | LGram | $O(n)^{*}$ | $O(g + n_{max})$ |
| lyndongrammar [6] | LGram | RLBWT | $O(n)$ | $\Omega(g)^{\P}$ |
| PFP [7] | Text | PFP | $O(n)$ | $O(\lvert D_1\rvert)$ |
| big-bwt [7] | PFP | RLBWT | $O(n)$ | $O(\lvert D_1\rvert + \lvert P_1\rvert)$ |
| r-pfbwt [8] | PFP | RPFP | $O(\lvert P_1\rvert)$ | $O(\lvert D_2\rvert)$ |
| r-pfbwt [8] | RPFP | RLBWT | $O(n)$ | $O(\lvert D_1\rvert + \lvert D_2\rvert + \lvert P_2\rvert)$ |

$n$: length of the text, $n_{max}$: length of the longest string in the text,$r$: number of runs in the BWT, $N$: number of strings in the text, $\kappa$: # of insert-heads, $\mathcal{R}$: reference string provided to CMS-BWT, $g$: size of the Lyndon SLP of the text, $D_1$: the dictionary of the PFP of the text, $P_1$: the parse of the PFP of the text, $D_2$: the dictionary of the PFP of $P_1$, $P_2$: the parse of the PFP of $P_1$

$^{*}$ expected time complexity

$^{\dagger}$ grlbwt is faster and smaller in practice than many other methods despite its worse time and space complexities

$^{\ddagger}$ requires providing a "reference" string that is similar to strings in the text

$^{\S}$ for texts where $\frac{n}{r} \geq \log n$, otherwise $O(n + r \log r)$

$^{\P}$ space complexity not explicitly stated in paper, RLBWT construction complexity used as lower bound

Table S2: Time and Memory for RLBWT construction on mtb152 and chr19.1000

| Tool | Threads | mtb152 Time (sec) | mtb152 Memory (MiB) | chr19.1000 Time (min) | chr19.1000 Memory (GiB) |
|---|---|---|---|---|---|
| CMS-BWT [3] | 1 | 24.66 | 666.4 | 51.55 | 24.5 |
| grlbwt [2] | 1 | 117.12 | 78.9 | 215.00 | 2.1 |
| | 2 | 69.26 | 228.7 | 109.93 | 16.0 |
| | 4 | 46.62 | 260.5 | 70.12 | 16.0 |
| | 8 | 35.45 | 324.7 | 48.29 | 16.1 |
| | 16 | 48.33 | 451.8 | 37.34 | 16.2 |
| | 32 | 57.26 | 745.9 | 32.98 | 16.4 |
| | 64 | 63.33 | 1,464.4 | 33.66 | 16.9 |
| lyndongrammar [6] | 1 | 44.43 | 168.2 | 78.82 | 8.6 |
| | 2 | 33.64 | 270.7 | 47.53 | 7.8 |
| | 4 | 22.35 | 270.1 | 30.26 | 7.7 |
| | 8 | 16.74 | 266.8 | 20.34 | 7.7 |
| | **16** | **13.98** | **260.4** | **15.27** | **8.4** |
| | 32 | 15.29 | 272.2 | 12.23 | 9.8 |
| | 64 | 16.91 | 408.1 | 11.56 | 12.6 |
| r-pfbwt [8] | 1 | 51.24 | 449.8 | $\geq$ 64.70* | $\geq$ 14.3* |
| | 2 | 49.37 | 449.8 | $\geq$ 64.74* | $\geq$ 14.3* |
| | 4 | 47.34 | 449.8 | $\geq$ 64.51* | $\geq$ 14.3* |
| | 8 | 46.64 | 449.8 | $\geq$ 64.47* | $\geq$ 14.3* |
| | 16 | 45.89 | 449.8 | $\geq$ 64.99* | $\geq$ 14.3* |
| | 32 | 46.51 | 449.8 | $\geq$ 65.12* | $\geq$ 14.3* |
| | 64 | 45.78 | 449.8 | $\geq$ 64.80* | $\geq$ 14.3* |

Bolded results are used for other experiments (as precursors to rlbwt2lcp and TeraLCP). ropebwt3 [4] was not tested because it has a parameter `‘‘-m batch size’’` that significantly affects run time and peak memory usage, and selecting an appropriate balance for small datasets is nontrivial. r-comp [5] was not tested because its implementation doesn't natively support string collections (and characters in the input text are bytes, limiting it to string collections with $\leq$ 256 strings. big-bwt [7] is not tested because it outputs the BWT instead of the RLBWT and r-pfbwt [8] is the algorithmic successor to big-bwt.

*: r-pfbwt crashed on all chr19.1000 runs (segmentation fault).

Table S3: Computing environments used for benchmarks in this paper

| location | name | GHz | Cores | CPU | RAM |
|---|---|---|---|---|---|
| University of Central Florida | ec326 | 1.8 | 48 | Intel(R) Xeon(R) Gold 5418N | 3 TiB |
| University of Central Florida | c0-5 | 2.2 | 128 | Intel(R) Xeon(R) Gold 6338N | 1 TiB |
| University of Central Florida | c0-4 | 2.1 | 64 | Intel(R) Xeon(R) CPU E5-2683 v4 | 0.5 TiB |
| Amazon Web Services | aws | 3.5 | 96 | x2iedn.24xlarge* | 3 TiB |

* Not the CPU, but type of rented AWS instance.

Table S4: Dataset Statistics

| | $n$ | $N$ | $r$ | $\frac{n}{r}$ |
|---|---|---|---|---|
| mtb152 | 1,341,467,444 | 304 | 6,740,559 | 199.0 |
| chr19.1000 | 118,250,232,020 | 2,000 | 91,081,449 | 1298.3 |
| human100 | 603,119,813,674 | 77,274 | 4,260,100,926 | 141.6 |
| human472 | 2,842,375,961,968 | 78,264 | 5,259,024,318 | 540.5 |
| CommonBacteria | 14,653,267,888,712 | 556,844,270 | 17,683,439,751 | 828.6 |

Datasets used in benchmarks throughout the paper. Each dataset, consists of the forward and reverse complement strands of the original dataset. mtb152, human100, and CommonBacteria were obtained from `https://doi.org/10.5281/zenodo.13147120`. human472 was obtained from `https://doi.org/10.5281/zenodo.11533210` [4]. chr19.1000 is chromosome 19 of 1,000 haplotypes generated as in [9]: "using the bcftools consensus tool to integrate variant calls from the phase-3 callset into chromosome 19 of the GRCh37 reference." $n$ is number of characters in the text. $N$ is number of strings in the text (counting forwards and reverse complements as distinct strings). $r$ is number of runs in the BWT.

Table S5: Run time and memory of TeraLCP for various datasets and threads

| | mtb152 | | chr19.1000 | | human100 | | human472 | | CommonBacteria | |
|---|---|---|---|---|---|---|---|---|---|---|
| threads | Sec | MiB | Hours | GiB | Hours | GiB | Hours | GiB | Hours | GiB |
| 1 | 247.07 | 159.24 | 16.10 | 2.79 | - | - | - | - | - | - |
| 2 | 122.83 | 158.75 | 8.01 | 2.79 | - | - | - | - | - | - |
| 4 | 64.24 | 159.26 | 4.53 | 2.79 | >24 | - | - | - | - | - |
| 8 | 34.99 | 157.98 | 2.34 | 2.79 | 20.88 | 136.10 | - | - | - | - |
| 16 | 19.28 | 159.31 | 1.22 | 2.79 | 10.74 | 136.18 | >24 | - | - | - |
| 32 | 10.82 | 158.75 | 0.63 | 2.79 | 5.66 | 136.32 | 22.86 | 171.66 | - | - |
| 64 | 7.27 | 159.53 | 0.35 | 2.79 | 3.82 | 136.56 | 12.69 | 171.83 | - | - |
| 128 | 5.88 | 159.39 | 0.21 | 2.80 | 2.23 | 136.99 | 8.44 | 172.16 | 48.53 | 613.65 |

Except for CommonBacteria, runs that took $>$ 24 hours were terminated early (denoted by ">24"). Runs with "-" in the hours column were not attempted due to the assumption they would take more than 24 hours. See a plot of this data in Figure S1.

Table S6: Memory usage (GiB) of LCP summary tools on random subsets of datasets

| | mtb152 | | | chr19.1000 | | | human100 | | |
|---|---|---|---|---|---|---|---|---|---|
| % of original | rb2lcp | pfpth | tera | rb2lcp | pfpth | tera | rb2lcp* | pfpth | tera |
| 1.56 | 0.11 | 0.15 | 0.12 | 2.15 | 3.22 | 2.13 | 46.95 | 144.37 | 78.08 |
| 3.13 | 0.12 | 0.17 | 0.12 | 3.38 | 4.14 | 2.15 | 61.39 | 196.96 | 95.24 |
| 6.25 | 0.15 | 0.19 | 0.15 | 5.72 | 5.60 | 2.15 | 79.08 | 255.52 | 107.43 |
| 12.50 | 0.20 | 0.23 | 0.16 | 10.37 | 8.25 | 2.20 | 105.82 | 320.70 | 112.63 |
| 25.00 | 0.30 | 0.29 | 0.16 | 19.76 | 13.18 | 2.31 | 154.98 | 411.95 | 123.64 |
| 50.00 | 0.51 | 0.44 | 0.17 | 38.57 | 22.58 | 2.50 | 257.05 | 554.87 | 127.90 |
| 100.00 | 0.92 | 0.70 | 0.17 | 76.00 | 40.66 | 2.83 | 460.18 | 791.92 | 137.01 |

Sequential grlbwt always used less memory than all LCP summary tools. rb2lcp is rlbwt2lcp, pfpth is pfp-thresholds, and tera is TeraLCP.
* Memory usage of rlbwt2lcp on human100 datasets estimated according to formula on its github.

Table S7: Min LCP Computation

| dataset | threads | tool | BWT time | BWT mem | Min LCP time | Min LCP mem | Total time | Total mem |
|---|---|---|---|---|---|---|---|---|
| mtb152 | 1 | CMS-BWT+rlbwt2lcp | 25 | 0.651 | 7:10 | 0.897 | 7:35 | 0.897 |
| | | pfp-thresholds | - | - | 2:44 | 0.698 | 2:44 | 0.698 |
| | | grlbwt+TeraLCP | 1:57 | 0.077 | 5:42 | 0.156 | 7:39 | 0.156 |
| | | lg+TeraLCP | 0:44 | 0.164 | 5:42 | 0.156 | 6:26 | 0.164 |
| | 128 | lg+rlbwt2lcp | 0:14 | 0.254 | 7:10 | 0.897 | 7:24 | 0.897 |
| | | pfp-thresholds | - | - | 2:18 | 0.698 | 2:18 | 0.698 |
| | | grlbwt+TeraLCP | 1:57 | 0.077 | 0:08 | 0.156 | 2:05 | 0.156 |
| | | lg+TeraLCP | 0:14 | 0.254 | 0:08 | 0.156 | 0:22 | 0.254 |
| chr19.1000 | 1 | CMS-BWT+rlbwt2lcp | 51:33 | 24.479 | 15:16:17 | 75.996 | 16:07:50 | 75.996 |
| | | pfp-thresholds | - | - | 3:29:38 | 40.656 | 3:29:38 | 40.656 |
| | | grlbwt+TeraLCP | 3:35:00 | 2.076 | 21:04:15 | 2.791 | 24:39:15 | 2.791 |
| | | lg+TeraLCP | 1:18:49 | 8.577 | 21:04:15 | 2.791 | 22:23:04 | 2.791 |
| | 128 | lg+rlbwt2lcp | 15:16 | 8.399 | 15:16:17 | 75.996 | 15:31:33 | 75.996 |
| | | pfp-thresholds | - | - | 2:52:14 | 40.655 | 2:52:14 | 40.655 |
| | | grlbwt+TeraLCP | 3:35:00 | 2.076 | 16:42 | 2.794 | 3:51:42 | 2.794 |
| | | lg+TeraLCP | 15:16 | 8.399 | 16:42 | 2.794 | 31:58 | 8.4 |

lg is lyndongrammar. When 128 threads are available, lyndongrammar is run with 16 threads. grlbwt and rlbwt2lcp are always run with exactly one thread. lyndongrammar in conjunction with TeraLCP always uses less memory and time than pfp-thresholds and any RLBWT construction tool with rlbwt2lcp.

#### S4 Supplementary Pseudocodes

**Algorithm 1:** IntervalPermutationToIntervalSequence($p, \pi$)

```
for i = 1 → r do
    I[i].first = p_i;
    π^{-1}[π[i]] = i;
q = 1;
for i = 1 → r do
    I[i].second = q;
    q = q + (p_{π^{-1}[i]+1} − p_{π^{-1}[i]});
return I;
```

**Algorithm 2:** InvertIntervalPermutation($p, \pi$)

```
for i = 1 → r do  π^{-1}[π[i]] = i ;
q_1 = 1;
for i = 1 → r do  q_{i+1} = q_i + (p_{π^{-1}[i]+1} − p_{π^{-1}[i]}) ;
return (q, π^{-1});
```

**Algorithm 3:** LFIntervalPermutation(RLBWT)

```
for c = 1 → σ + 1 do  C_r[c] = 0 ;
/* Step 1: compute run count per character, shifted by 1 to simplify exclusive
   prefix sum                                                                  */
for i = 1 → r do  C_r[RLBWT[i].c + 1] = C_r[RLBWT[i].c + 1] + 1 ;
// Step 2: exclusive prefix sum
for C_r[c] = 2 → σ + 1 do  C_r[c] = C_r[c − 1] + C_r[c] ;
// Step 3: compute p and π
p' = 1;
for i = 1 → r do
    p[i] = p';
    p' = p' + RLBWT[i].len;
    C_r[RLBWT[i].c] = C_r[RLBWT[i].c] + 1;
    π[i] = C_r[RLBWT[i].c];
return (p, π);
```

**Algorithm 4:** splitIntervalPermutation(RF, $D_{pair}, (p_\psi, \pi_\psi)$)

```
// Count output intervals before
index = 1;
for i = 1 → r do
    splitInto=0;
    while index ≤ |D_pair| and D_pair[index].first < p_ψ[i + 1] do
        index = index + 1;
        splitInto = splitInto + 1;
    outputSplitInto[π[i] + 1] = splitInto;
outputSplitInto[1] = 1;
for i = 2 → r do  outputSplitInto[i] = outputSplitInto[i − 1] + outputSplitInto[i] ;
// Split disjoint interval permutation
index = 1;
for i = 1 → r do
    interval = splitInto[π_ψ[i]];
    while index < |D_pair| and D_pair[index + 1].first < p_ψ[i + 1] do
        RF′[index] = (RF[i].c, D_pair[index + 1].first − D_pair[index].first);
        π′[index] = interval;
        interval = interval + 1;
        index = index + 1;
    RF′[index] = (RF[i].c, p_ψ[i + 1] − D_pair[index].first);
    π′[index] = interval;
    interval = interval + 1;
    index = index + 1;
return RF′, π′;
```

**Algorithm 5:** getPsiCoordLastSuffix(RF)

```
N = 0;
for i = 1 → |RF| do
    if RF[i].c = $_* then                    // if RF[i].c is a terminal character
        N = N + 1;
return (N, N, 1);
```

**Algorithm 6:** PhiIntervalPermutation(RLBWT, $\mathcal{M}_\psi$)

$(p_\psi, \pi_\psi) =$ InvertIntervalPermutation(LFIntervalPermutation(RLBWT)); `// Algorithms 2 and 3`
`// Step 1: Compute RF'`
**for** $i = 1 \to r$ **do** $\mathrm{RF}[i] = \mathrm{RLBWT}[\pi_\psi[i]]$ ;
$\mathrm{RF}', \pi'_\psi =$ splitIntervalPermutation(RF, $\mathcal{M}_\psi.D_{pair}$, $\pi_\psi$); `// Algorithm 4`
`// Step 2: Compute` $p$ `and` $SA_{inp}$
$begin = crd = \mathcal{M}_\psi$(getPsiCoordLastSuffix(RF$'$)); `// Algorithm 5`
$tops = 1, s = 1, phiIntervalBegin = 1$;
**repeat**
  **if** $crd.off = 1$ **then**
    **if** $crd.interval = 1$ *or* $RF'[crd.interval].c \neq RF'[crd.interval - 1].c$ *or* $\pi'_\psi[crd.interval] \neq \pi'_\psi[crd.interval - 1] + 1$ **then**
      $SA_{inp}[crd.interval] = tops$;
      $p[tops] = phiIntervalBegin$;
      $tops = tops + 1$;
      $phiIntervalBegin = s + 1$;
  $s = s + 1$;
  $crd = \mathcal{M}_\psi(crd)$;
**until** $crd = begin$;
`// Step 3: Compute` $\pi$
$bots = 1, s = 1$;
$crd = begin$;
**repeat**
  **if** $crd.pos = RF'[crd.interval].len$ **then**
    **if** $crd.interval = |RF'|$ *or* $RF'[crd.interval].c \neq RF'[crd.interval + 1].c$ *or* $\pi'_\psi[crd.interval] \neq \pi'_\psi[crd.interval + 1] - 1$ **then**
      $next = (crd.interval \mod |RF'|) + 1$;
      $\pi[SA_{inp}[next]] = bots$;
      $bots = bots + 1$;
  $s = s + 1$;
  $crd = \mathcal{M}_\psi(crd)$;
**until** $crd = begin$;
return $(p, \pi)$;

**Algorithm 7:** NaiveiPLCP(RLBWT, $\mathcal{M}_\psi$)

```
(p_ψ, π_ψ) = InvertIntervalPermutation(LFIntervalPermutation(RLBWT));  // Algorithms 2 and 3
// Step 1:  Compute RF′
for i = 1 → r do RF[i] = RLBWT[π_ψ[i]] ;
RF′, π′_ψ = splitIntervalPermutation(RF, M_ψ.D_pair, π_ψ);                      // Algorithm 4
// Get SA_inp from PhiIntervalPermutation algorithm as well as return values
(p_φ, π_φ, SA_inp) = PhiIntervalPermutation(RLBWT, M_ψ);                         // Algorithm 6
for i = 1 → |RF′| do
    if i ≠ r and RF′[i] = RF′[nextRun] and π′_ψ[nextRun] = π′_ψ[i] + 1 then
        continue;
    nextRun = (i mod |RF′|) + 1;
    crd_a = (p_ψ[i + 1] − 1, i, p_ψ[i + 1] − p_i);
    crd = (p_ψ[nextRun], nextRun, 1);
    len = 0;
    while RF′[crd_a.interval] = RF′[crd.interval] do
        crd_a.interval = M_ψ(crd_a.interval);
        crd.interval = M_ψ(crd.interval);
        len = len + 1;
    iPLCP[SA_inp[nextRun]] = p_φ[SA_inp[nextRun] + 1] − p_φ[SA_inp[nextRun]] − 1 + len;
return iPLCP;
```

**Algorithm 8:** LineariPLCP(RLBWT, $\mathcal{M}_\psi$)

```
(p_ψ, π_ψ) = InvertIntervalPermutation(LFIntervalPermutation(RLBWT));  // Algorithms 2 and 3
// Step 1: Compute RF′
for i = 1 → r do RF[i] = RLBWT[π_ψ[i]] ;
RF′, π′_ψ = splitIntervalPermutation(RF, M_ψ.D_pair, π_ψ);  // Algorithm 4
// Get SA_inp from PhiIntervalPermutation algorithm as well as return values
(p_φ, π_φ, SA_inp) = PhiIntervalPermutation(RLBWT, M_ψ);  // Algorithm 6
I_φ = IntervalPermutationToIntervalSequence(p_φ, π_φ);  // Algorithm 1
for i = 1 → r do SA_inp^{-1}[SA_inp[i]] = i ;
// Step 2: Get ISA samples
crd = M_ψ(getPsiCoordLastSuffix(RF′));  // Algorithm 5
sampleInterval = ⌊n/r⌋;
for s = 1 → n do
    if s mod sampleInterval = 0 then ISA_r[s/sampleInterval] = crd ;
    crd = M_ψ(crd);
// Step 3: Compute PLCP values
crd = M_ψ(getPsiCoordLastSuffix(RF′));  // Algorithm 5
s = 1;
for i = 1 → r do
    // Forward crd if it is behind
    while s < p_φ[i + 1] − 1 do
        s = s + 1;
        crd = M_ψ(crd);
    // Compute crd_a in O(n/r) time
    s_a = I_φ[i].second + (s − I_φ[i].first);
    s′_a = ⌊I_φ[i].second / sampleInterval⌋;
    crd_a = ISA_r[s′_a];
    s′_a = s′_a × sampleInterval;
    while s′_a ≠ s_a do
        crd_a = M_ψ(crd_a);
        s′_a = s′_a + 1;
    // Compute PLCP value
    while RF′[crd.interval] = RF′[crd_a.interval] do
        s = s + 1;
        crd = M_ψ(crd);
        crd_a = M_ψ(crd_a);
    iPLCP[i] = s − I_φ[i].first;
return iPLCP;
```
